# Vulnerability of triple negative breast cancer to daraxonrasib pan-RAS(on) inhibition

**DOI:** 10.64898/2026.09.16.752096

**Authors:** Michael P. East, J. Felix Olivares-Quintero, Denis O. Okumu, Kevin R. Mott, Robert W. Sprung, Austin A. Whitman, Chinmaya U. Joisa, Xin Chen, Qiang Zhang, Petra Erdmann-Gilmore, Yiling Mi, James P. Malone, Sonam Bhatia, Ian C. McCabe, Yi Xu, Matthew D. Sutcliffe, Patricia A. Spears, Jon S. Zawistowski, Charles M. Perou, H. Shelton Earp, Lisa A. Carey, Jen Jen Yeh, David L. Spector, Daniel Stover, Shawn M. Gomez, Philip M. Spanheimer, R. Reid Townsend, Gary L. Johnson

**Affiliations:** Department of Pharmacology, University of North Carolina at Chapel Hill, Chapel Hill, NC, USA; Lineberger Comprehensive Cancer Center, University of North Carolina at Chapel Hill, Chapel Hill, NC, USA; Division of Endocrinology, Metabolism, and Lipid Research, Department of Medicine, Washington University School of Medicine, St. Louis, MO, USA; Joint Department of Biomedical Engineering, University of North Carolina at Chapel Hill, Chapel Hill, NC, USA; Department of Pharmaceutical Clinical Sciences, Campbell University, Buies Creek, NC, USA; Cold Spring Harbor Laboratory, Cold Spring Harbor, New York, NY, USA; Department of Genetics, University of North Carolina at Chapel Hill, Chapel Hill, NC, USA; Department of Medicine, University of North Carolina at Chapel Hill, Chapel Hill, NC, USA; Department of Surgery, University of North Carolina, Chapel Hill, NC, USA; The Ohio State University Comprehensive Cancer Center Arthur G James Cancer Hospital and Richard J Solove Research Institute, Columbus, Ohio

**Keywords:** Triple negative breast cancer (TNBC), adaptive resistance, kinome reprogramming, RAS, daraxonrasib, pan-RAS(on) inhibitor, trametinib, RAS-RAF-MEK-ERK pathway inhibition

## Abstract

Triple-Negative Breast Cancer (TNBC) is a clinical subtype where aberrant expression of receptor tyrosine kinases (RTKs) leads to the activation of the RAS-RAF-MEK-ERK pathway contributing to tumor progression. Therapies targeting components of the RAF-MEK-ERK pathway in TNBC have been disappointing due to rapid onset of adaptive reprogramming of the RTK landscape leading to pathway reactivation. To overcome the dynamic heterogeneity of RTK expression and their convergence in activating RAS, treatment with the pan-RAS(on) inhibitor daraxonrasib (RMC-6236), which inhibits RAS-GTP regulation of RAF kinases, inhibited proliferation independent of TNBC kinotype or oncogenic RAS mutation in three of four TNBC cell lines *in vitro* and two patient-derived TNBC xenografts in mice. Daraxonrasib in combination with the MEK inhibitor trametinib prevented reactivation of the RAF-MEK-ERK pathway observed with each inhibitor alone and further enhanced inhibition of tumor growth in TNBC cell lines and xenografts. Inhibiting pan-RAS(on) RAS-GTP with daraxonrasib, alone or in combination with a MEK inhibitor, provides a new strategy for the treatment of TNBC that inhibits RTK activation of the RAS-RAF-MEK-ERK pathway and tumor proliferation.

## Introduction

Breast cancer patients lacking estrogen receptor (ER), progesterone receptor (PR) and the receptor tyrosine kinase ERBB2/HER2 are referred to as Triple Negative Breast Cancers (TNBC). Until recently chemotherapy has been the standard treatment for TNBC ^1^, but the immune checkpoint inhibitor pembrolizumab is now also approved for use in early stage and metastatic TNBCs in combination with chemotherapy^2, 3^. In addition, the antibody-drug-conjugates (ADC) sacituzumab govitecan, datopotamab deruxtecan, and trastuzumab deruxtecan have shown efficacy in patients with metastatic TNBC. Even with these advances in treatment, TNBC patients still experience the worst survival outcomes compared to ER+/HER2- and HER2+ patients. It is therefore imperative that the vulnerabilities of TNBC be further characterized for developing new therapeutic strategies for treatment. Genomic studies have shown that 22% of TNBC are EGFR amplified, 31% are KRAS amplified, and 32% are BRAF amplified resulting in an enhanced activation of the RAS-RAF-MEK-ERK signaling pathway ^4^. Inhibition of the RAS-RAF-MEK-ERK pathway in TNBC slows tumor growth initially but inhibiting one component of the pathway, such as trametinib inhibition of MEK, is not durable ^5, 6, 7, 8^. Epigenetic and transcriptomic changes resulting from MEK inhibition rapidly alter the expression of protein kinases including multiple receptor tyrosine kinases (RTKs), which activate Raf-MEK-ERK in a RAS dependent signaling network resulting in an adaptive resistance overcoming growth arrest ^5, 6, 7, 9, 10^. Mechanistically with MEK-ERK inhibition, cMYC is rapidly degraded resulting in chromatin remodeling and transcriptional changes of tyrosine and serine/threonine protein kinases^5, 6^. Similar inhibitor-induced adaptive proteomic reprogramming is seen with many RTK and serine/threonine kinase inhibitors as well as RAS inhibitors studied in different cancers^11, 12, 13^.

As a result, a major challenge for developing new targeted therapies for TNBC is the inter-tumoral heterogeneity of the expressed kinome at baseline and in response to therapy. We have used a human kinase parallel reaction monitoring (PRM) peptide library coupled with SureQuant triggered acquisition mass spectrometry to determine the dynamic variability of the breast cancer kinome^14, 15^. SureQuant/PRM analysis quantitated an overlapping heterogenous expression of RTKs across primary tumors, organoids and cell lines. Similar findings were observed at the transcriptional level by RNAseq in patient samples from two TNBC clinical trials investigating the action of the MEK inhibitor trametinib ^6, 8^. Given the challenges in targeting RTKs due to this array of RTK expression in TNBC and that RTKs converge on RAS to activate RAF-MEK-ERK signaling, we hypothesized that effective chemical inhibition of RAS would more effectively reduce TNBC growth by targeting the convergence point of heterogenous activation. To do this, we tested the novel pan-RAS(on) inhibitor daraxonrasib (RMC-6236) that inhibits both wild-type RAS and oncogenic RAS mutants^13, 16^. Daraxonrasib inhibited TNBC cell growth *in vitro* and increased survival *in vivo* with patient xenografts treated in NSG mice. However, adaptive reprogramming and pathway reactivation similar to that seen with MEK inhibition was observed. Combining daraxonrasib with trametinib enhanced efficacy compared to either drug alone providing a pharmacological method to overcome the dynamic expression and adaptive reprogramming of RTKs.

## Results

### Breast cancer patient tumors and preclinical models exhibit heterogenous protein kinase kinotypes

Proteomic analysis of the kinome from breast cancer patient samples, cell lines, and patient derived organoids (PDOs) was performed using a targeted parallel reaction monitoring (PRM) proteomics approach with heavy amino acid labeled reference peptides spiked into each sample at known concentrations^17, 18^. Stable isotope labeled (SIL) peptides were uniformly ^13^C and ^15^N labeled at C-terminal amino acids and were used to trigger acquisition of endogenous peptide data using Thermo SureQuant software to simultaneously measure abundance of both peptides for accurate quantitation of kinase abundance^18^. Kinotypes of 9 TNBC (HCC70, HCC1143, HCC1806, Hs578T, MDA-MB-231, MDA-MB-468, SUM102PT, SUM149PT, and SUM159PT) and two HER2+ cell lines (SKBR3, BT474), three TNBC PDOs (NH85TSc, NH87TT, and NH95TT), and eight TNBC patient tumors were analyzed by SureQuant/PRM. Variance of kinase protein expression in cell lines, organoids and patient tumors is summarized in kinome trees in Figure 1A. Circle size represents variance of expression with increasing circle size indicating a greater variance across the cell lines, organoids or patient tumors for that specific kinase. The kinome trees for each of the breast cancer models used in this study demonstrate diversity of kinase expression in each subfamily of the kinome. The greatest variance is seen in the RTK/TK subfamily of kinases, which is consistent across models. Dot blots in Figure 1B (cell lines), 1C (PDOs) and 1D (patient tumors) show expression of 16, 22 and 24 RTKs, respectively. Patient tumors have a microenvironment component that likely contributed to the expression of some kinases, but the organoids and cell lines are free of stromal cells. This analysis shows that multiple RTKs and TKs are expressed in TNBC at varying levels. RTKs mechanistically activate RAS and the RAS-MEK-ERK signaling pathway is a primary proliferative pathway in TNBC^4, 5, 6^ (Figure 1E). Thus, dynamic variation in RTK expression represents an important clinical challenge as it limits the utility of RTK inhibitors, which consist of approximately half of all kinase inhibitors approved by the FDA (Figure 1F).

**Figure 1:**
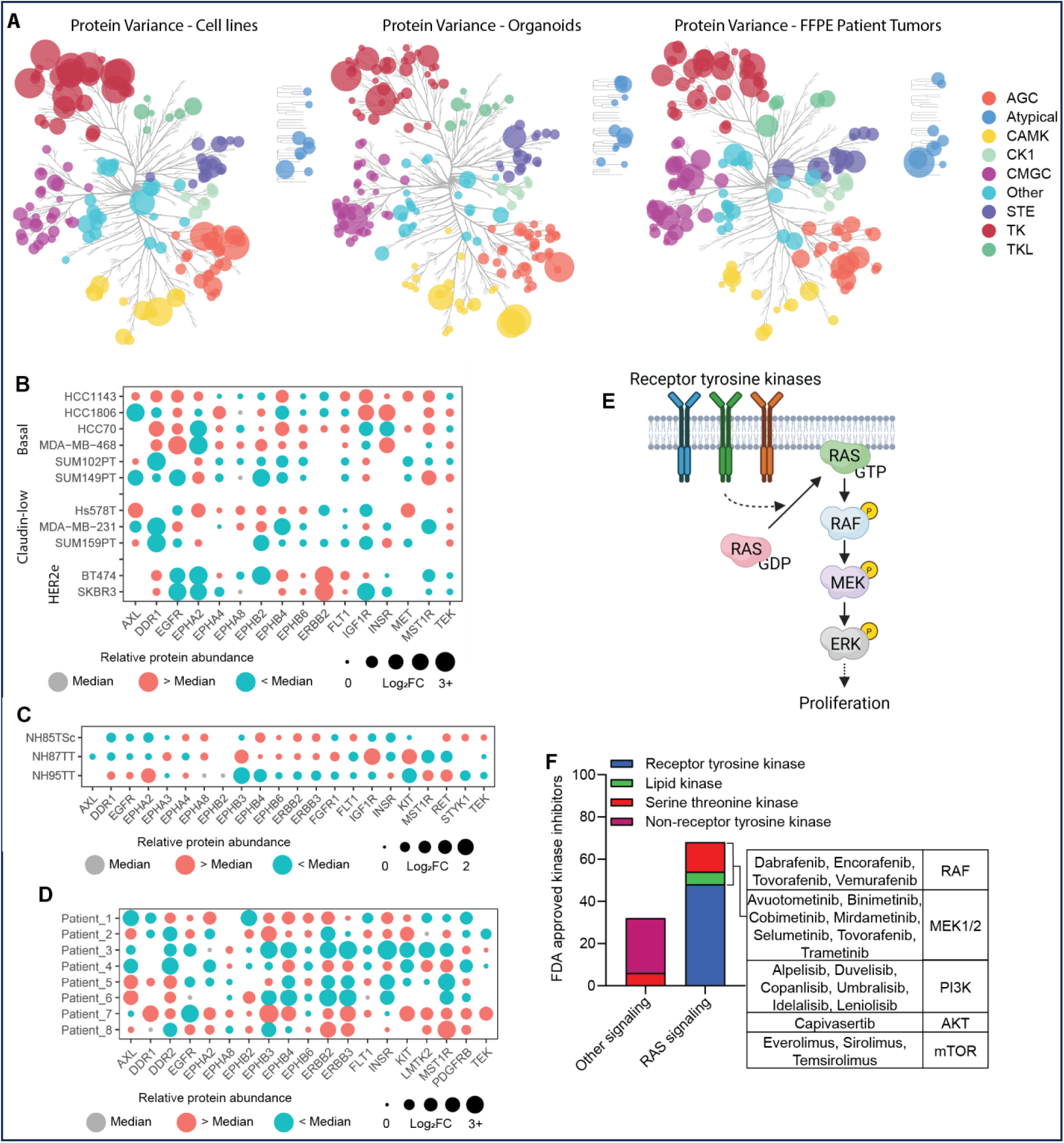
Receptor tyrosine kinases are the most variant kinases across breast cancer preclinical models and patient tumor kinotypes. **A)** Variance in receptor tyrosine kinase (RTK) protein abundance determined using internal standard triggered targeted proteomics (SureQuant/PRM) was calculated for each kinase across cell lines, patient derived organoids (PDOs), and flash-frozen patient tumor samples separately. Circle sizes correlate with relative variance levels and circle colors indicate the subfamily of each kinase. **B-D)** RTK expression heterogeneity in cell lines (**B**), PDOs (**C**), formalin fixed paraffin embedded patient tumors (**D**), and flash-frozen patient tumors (**E**). Protein expression levels were median centered and the difference from the median indicated by circle size and color following log_2_ transformation. Red circles indicate expression levels higher than the median and blue circles lower than the median. Gray circles indicate equivalence to the median. Shown are all the RTKs quantified from each sample type. Kinase expression levels from patient tumor samples were scaled to center the distribution of the data to zero after median normalization to account for differences in cellularity and quality across samples. **F)** Summary of on-label targets for all FDA-approved kinase inhibitors.

### Action of the pan-RAS(on) inhibitor daraxonrasib in TNBC

Rational treatment design based upon targeting one or two RTKs with RTK inhibitors is not feasible when considering larger TNBC patient cohorts. Thus, we hypothesized that targeting inhibition of RAS would inhibit the convergence point of heterogenous RTK activation of the RAS-RAF-MEK-ERK pathway. Daraxonrasib is a pan-RAS(on) inhibitor that binds RAS proteins having GTP bound versus inactive RAS having GDP bound. Daraxonrasib was developed by Revolution Medicine and is now FDA approved for treatment of pancreatic ductal adenocarcinoma^13, 16^. Daraxonrasib remodels the surface of the chaperone protein cyclophilin A (CypA) to generate a high affinity binding interface for the three different RAS proteins (KRAS, HRAS, NRAS) (Figure 2A). The CypA-daraxonrasib complex selectively binds activated RAS-GTP (RASon) but does not bind RAS-GDP and forms a stable, ternary complex that inhibits RAS signaling by preventing interactions with effector proteins including RAF kinases. Thus, daraxonrasib is a panRAS-GTP inhibitor that binds both mutant activated and wild-type RAS activated proteins^16^. Because daraxonrasib selectively binds RAS-GTP (pan-RAS(on), there is a pool of inactive RAS-GDP proteins that can be activated and daraxonrasib can titrate inhibition of the RAS proteins over time^13, 19^. Daraxonrasib as a panRAS(on) inhibitor has not been tested in TNBC and we now demonstrate its utility in inhibiting TNBC model systems both *in vitro* and *in vivo*.

**Figure 2:**
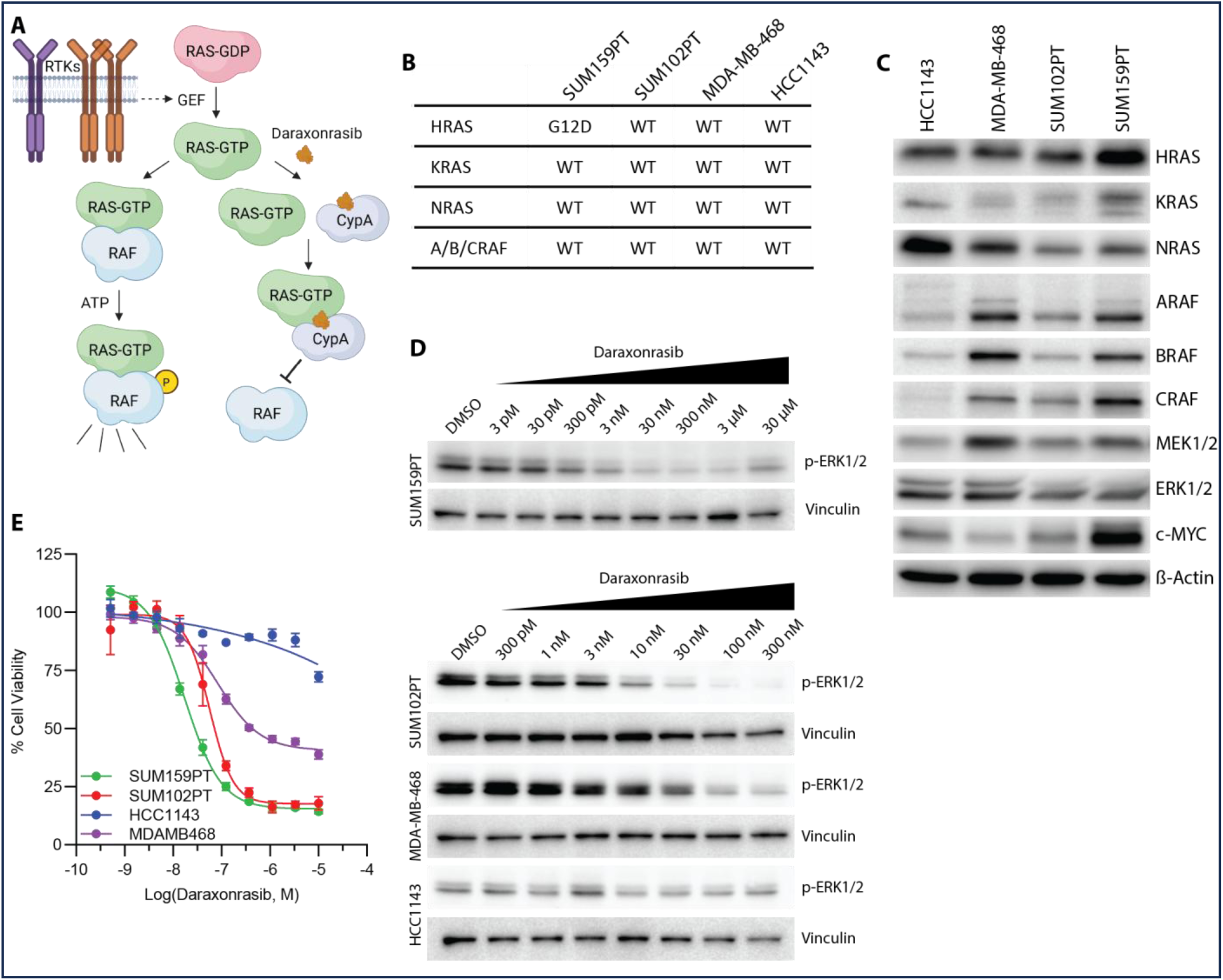
The pan-RAS-ON inhibitor daraxonrasib (RMC-6236) inhibits growth and MEK/ERK pathway activation in TNBC cell lines. **A)** RMC-6236 binds to cyclophilin A (CypA) and creates a high affinity interface for RAS-GTP binding. Stable ternary RAS-GTP/daraxonrasib/CypA complexes occlude binding and activation of RAS-GTP effectors including RAF kinases. RAS-GTP binding is stimulated by upstream RTKs. **B-C)** Molecular and proteomic characterization of four TNBC cell lines used to test the efficacy of daraxonrasib. Only SUM159PT cells harbored an oncogenic RAS mutation (**B**) but variable expression of other MEK/ERK pathway members was observed at the protein level (**C**). Actin was used as a loading control. **D)** Levels of activating phosphorylation of ERK1/2 were reduced upon treatment with daraxonrasib. Vinculin served as a loading control. Potency was highest in oncogenic RAS mutant SUM159PT cells. **E)** Daraxonrasib showed dose dependent growth inhibition in TNBC cell lines. Cells were treated with increasing concentrations of daraxonrasib for 72 hours and viability determined using Cell-Titer-Glo. Panel A was created in BioRender.

We evaluated daraxonrasib in a panel of four TNBC cell lines and, although RAS mutations are uncommon in TNBC, we included SUM159PT with an oncogenic HRAS (G12D) mutation. All other RAS and RAF proteins were wild-type (summarized in Figure 2B) but RAS-MEK-ERK pathway members ranged in protein expression levels (Figure 2C). Activating phosphorylation marks on ERK1/2 were used as readouts for RAS-RAF-MEK-ERK pathway inhibition by daraxonrasib (Figure 2D). SUM159PT, cells expressing an HRAS(G12D) mutation, were the most sensitive with a maximal effect observed at ∼30 nM daraxonrasib, the other cell lines reaching maximal loss of pERK signal at ∼100 nM. Daraxonrasib inhibited growth in three of the four cell lines in 72 hr growth assays with only a minor effect observed in HCC1143, where inhibition of the pERK signal was also less robust.

Studies of oncogenic RAS driven cancers have shown transcriptional plasticity in response to RAS inhibition similar to transcriptional effects observed with trametinib treatment ^10, 11, 12^. To investigate the effects of daraxonrasib on the transcriptome in patient tumors, we modified an operating room-to-laboratory pipeline^20, 21, 22^ to measure drug responses in primary human tumors. Needle biopsies were taken from three TNBC patients, rapidly minced, and cultured *in vitro* for 24 hours with 100 nM daraxonrasib or vehicle (DMSO) and then analyzed by RNAseq. Samples were never frozen and viability was preserved over treatment. The baseline expression levels of RTKs in the vehicle treated samples (Figure 3A) measured ∼50 RTKs expressed in each *ex vivo* tumor cell population with enrichment of distinct sets of RTKs in each *ex vivo* tumor. The heatmap in Figure 3A shows relative expression levels based on the median across the three *ex vivo* tumors. Treatment with daraxonrasib resulted in several thousand Differentially Expressed Genes (DEGs) per *ex vivo* tumor (Figure 3B), including RTKs (Figure3C), with distinct reprogramming of RTK kinotype in each *ex vivo* tumor (Figure 3D). The kinotype reprogramming included transcriptional changes in expression of the ligands required for activation of expressed RTKs (Suppl Fig. 1). Reprogramming of the transcriptome and RTK landscape in *ex vivo* TNBC patient tumors treated with daraxonrasib closely paralleled the effects of trametinib observed in clinical trials (Suppl Figure 2). To compare the effects of daraxonrasib and trametinib, we treated two TNBC cell lines with each inhibitor for 24 hours followed by RNAseq. Robust transcriptional reprogramming was observed in both cell lines in response to either daraxonrasib or trametinib (Figure 4A). Treatment with daraxonrasib resulted in 1,652 DEGs (≥ 2-fold change, p-adj < 0.05) in SUM159PT and 1,221 DEGs in SUM102PT cells, respectively. Similarly, trametinib treatment resulted in 2,688 and 2,827 DEGs in SUM159PT and SUM102PT cells. DEGs for each inhibitor in each cell line were largely overlapping (Figure 4B) with most of the daraxonrasib DEGs overlapping with trametinib DEGs. Comparing expression changes of each gene that changed in a statistically significant manner (p-adj < 0.05) in response to either inhibitor, there was a strong, linear correlation between daraxonrasib and trametinib in both cell lines (Figure 4C). Linear regression analysis yielded R^2^ values of 0.91 and 0.78 for SUM159PT and SUM102PT, respectively. Thus, transient inhibition of MEK-ERK signaling was sufficient to phenocopy inhibition of RAS, which can activate several signaling pathways in addition to RAF-MEK-ERK, suggesting that RAF-MEK-ERK signaling is the dominant RAS regulated pathway in TNBC similar to that seen with mutant RAS driven pancreatic ductal adenocarcinoma^23, 24^.

**Figure 3:**
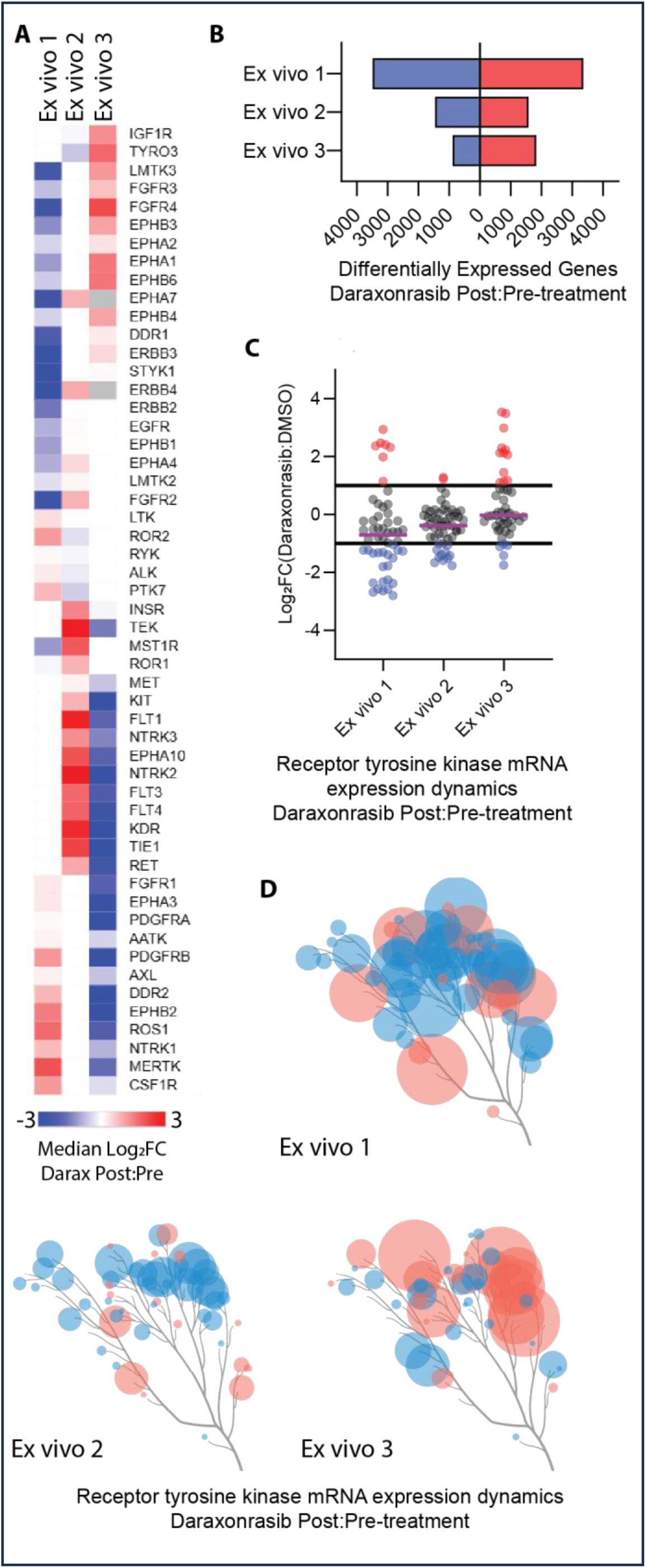
Daraxonrasib induces changes in the transcriptome in e*x vivo* treated TNBC patient tumors A-. **C)** Needle biopsies from TNBC patient tumors were collected and immediately minced and cultured in media containing vehicle alone (DMSO) or 100 nM RMC-6236 for 24 hours *in vitro*. Samples were then harvested and analyzed by RNAseq. **A)** RTK expression levels in DMSO treated samples were compared after median centering. Shading on the heatmap is based on the fold difference from the median with the darkest colors indicating an 8-fold or greater difference from the median expression level. Gray coloration indicates expression levels below 5 counts. **B)** The number of differentially expressed genes (DEGs) are summarized for each *ex vivo* patient sample. DEGs were defined as transcripts whose abundance changed by at least 2-fold in response to daraxonrasib treatment. Red and blue bars indicate transcripts that were up or down regulated by daraxonrasib treatment, respectively. **C-D)** Dynamic reprogramming of RTK transcripts is summarized for each patient. **D)** RTK kinome trees summarize the dynamic changes in RTK transcripts in response to daraxonrasib. Each circle indicates an RTK that was expressed in the indicated patient. Red and blue colors indicate up and down regulated transcripts, respectively. Circle size correlates with the magnitude of change with the largest circles indicating an 8-fold or larger change in expression. **E-G)**

**Figure 4:**
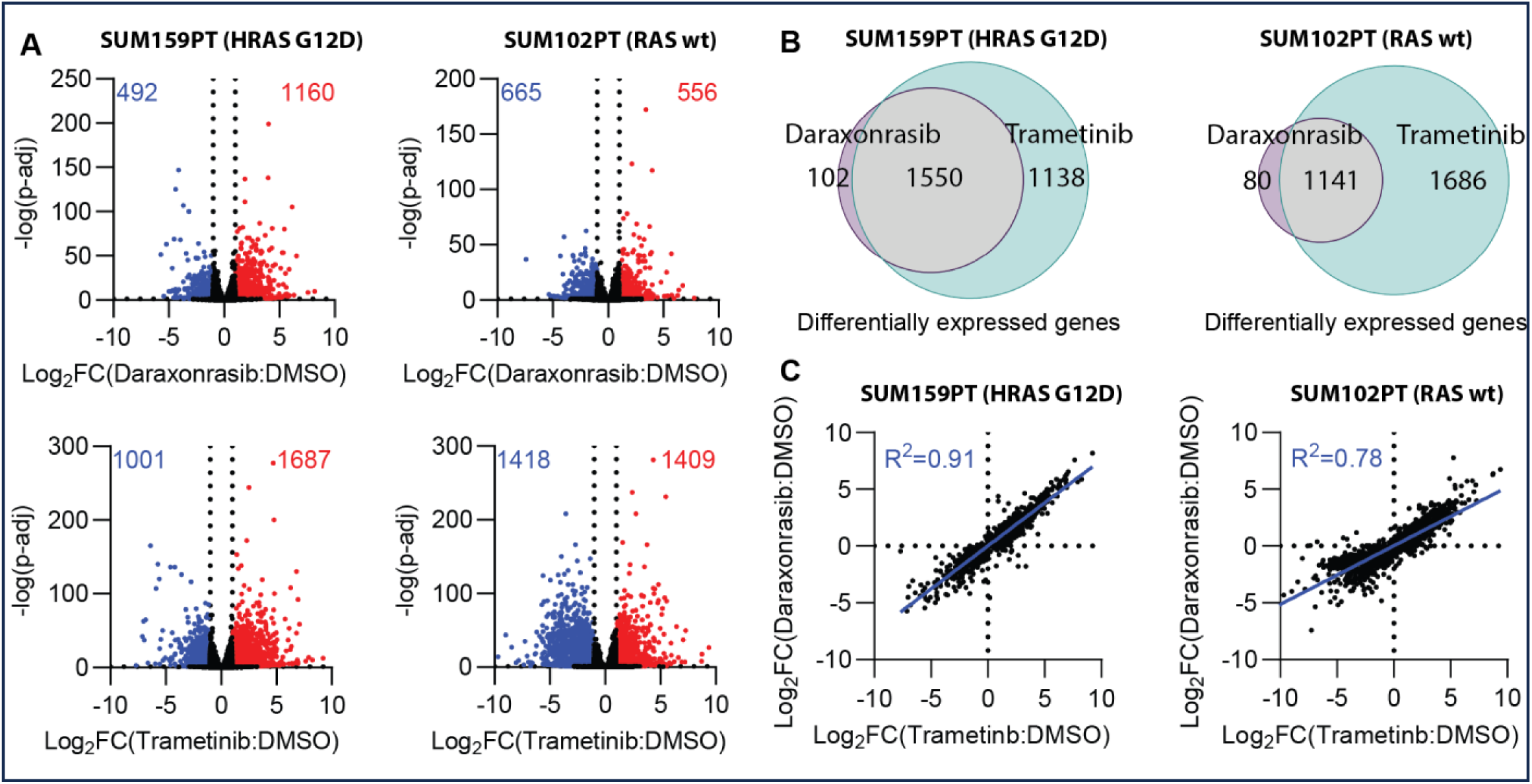
Daraxonrasib phenocopies trametinib induced transcriptional changes in TNBC cell lines. Two TNBC cell lines, one with an oncogenic HRAS mutation, SUM159PT, and another with WT RAS, SUM102PT, were treated with daraxonrasib or trametinib for 24 hours and transcriptional dynamics analyzed by RNAseq. **A)** Volcano plots demonstrate the number of DEGs (2-fold change in expression, p-adj < 0.05) in response to the indicated inhibitor in each cell line. **B)** The effects of daraxonrasib on the transcriptome were highly similar to trametinib in both TNBC cell lines. DEGs (≥ 2-fold, p-adj < 0.05) from either treatment are compared with Venn diagrams and are largely overlapping. **C)** Transcripts that were significantly altered (p-adj < 0.05) by either inhibitor are compared using scatter plots. The Log_2_ transformed fold changes in response to each inhibitor were plotted against each other and linear regression analysis performed resulting in R^2^ values of 0.91 and 0.78 for SUM159PT and SUM102PT, respectively.

### Pan-Ras(on) inhibition by daraxonrasib blocks adaptive resistance to trametinib-induced growth inhibition

The adaptive resistance to trametinib in TNBC is driven by transcriptional reprogramming and MEK-ERK pathway reactivation associated with the expression of RTKs^5, 6, 7^. Based on the hypothesis that daraxonrasib would inhibit the RTK-driven RAS(on) activation of the RAF-MEK-ERK pathway, we tested whether targeting RAS(on) with daraxonrasib would enhance trametinib inhibition of cell growth compared to single-agent trametinib. In short-term, 96-hour growth assays in three TNBC cell lines, combination with daraxonrasib greatly sensitized cells to trametinib treatment (Figure 5A). Increasing concentrations of daraxonrasib significantly left-shifted the trametinib response curves in a dose-dependent manner in each cell line tested, most likely because of suppression of RAF activation in response to RAS(on) inhibition. Neither compound alone or in combination induced cell death and, since cells weren’t killed, the response curves did not approach zero so the effective concentration 50 (EC50) of trametinib was calculated as the midpoint between the floor and ceiling of each response curve (Figure 5B). Based on the hypothesis that daraxonrasib would inhibit the RTK-driven activation of the RAF-MEK-ERK pathway, the RAS(on) inhibitor reduced the EC50 of trametinib by approximately 100-fold in HCC1143 and MDA-MB-468 cells and by approximately 10-fold in SUM102PT cells. To test the durability of growth inhibition by the daraxonrasib + trametinib combination, 14-day colony formation assays were performed on four TNBC cell lines including each of the lines tested at 96 hours and SUM159PT. At concentrations where each compound alone had only a minor effect on cell growth, the combination had a profound inhibition on cell growth (Figure 5C). Quantitation of cell growth from three biological replicates showed a greater than additive effect of the two compounds when used in combination (Figure 5D). A 2x3 combination matrix of drug concentrations was used to calculate Delta Bliss sum negative score for each cell line as a measure of *in vitro* synergy^25^. The DBsumNeg scores ranged from -1.5 to -2.6 indicating a synergistic relationship between the two compounds *in vitro*.

**Figure 5:**
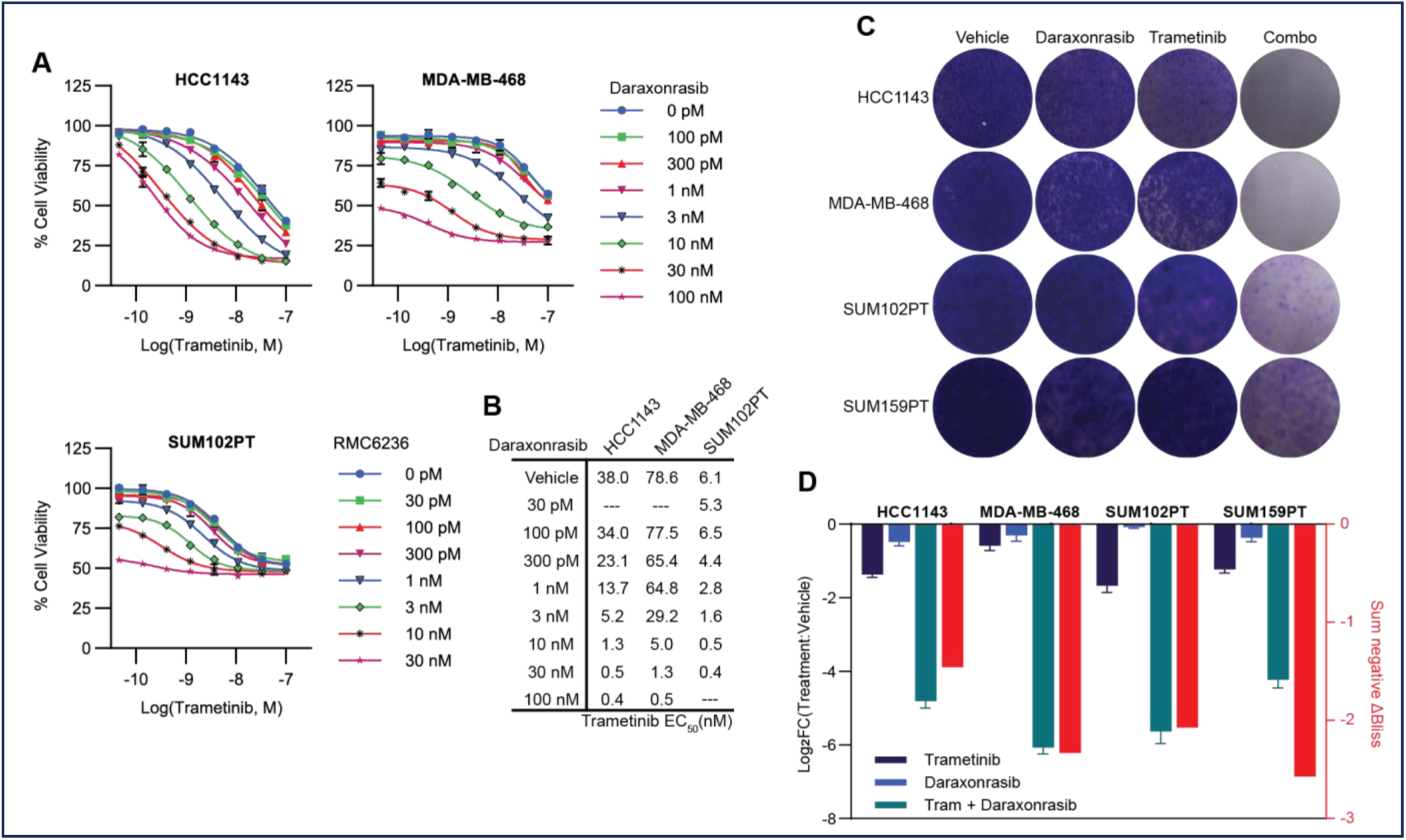
Pan-RAS(on) inhibition in combination with MEK-ERK inhibition inhibits TNBC cell growth and onset of adaptive resistance. **A)** Three TNBC cell lines were treated for 96-hours with trametinib alone or in combination with daroxonrasib across a range of doses of each compound. In each of the three cell lines, daraxonrasib sensitized cells to trametinib as indicated by the left-shift in the trametinib response curves. **B)** The EC50 of trametinib in the presence of each concentration of daraxonrasib was calculated based on the midpoint between the floor and ceiling of each trametinib response curve. Daraxonrasib shifts the EC50 of trametinib by 10 to 100-fold. **C)** 14-day colony formation assays show synergistic growth inhibition between trametinib and daraxonrasib in four TNBC cell lines. Trametinib was used at 3 nM for all cell lines and daraxonrasib was used at 1 nM (SUM102PT, SUM159PT) or 10 nM (HCC1143, MDA-MB-468). **D)** Quantification of the colony formation assay (**C**) across three biological replicates. Error bars indicate standard error of the mean. The sum negative ΔBliss score was calculated across 3 x 4 concentration grids as a quantitative measure of synergy and is indicated by the red bars.

Transcriptional reprogramming of the RTK landscape in response to trametinib drives adaptive resistance and reactivation of MEK-ERK signaling and RAS activation is an essential node in RTK activation of the RAF-MEK-ERK pathway. We used a RAS pulldown activation assay based on the RAS binding domain of cRAF to selectively precipitate activated RAS-GTP from lysates of tumor cells treated with trametinib for 0, 4, or 24 hours (Fig 6A). MDA-MB-468 is a TNBC cell line with wild-type RAS and SUM159PT is a TNBC harboring an oncogenic G12D HRAS mutation. Activated RAS pulldown revealed a time-dependent trametinib-induced increase in RAS activation in both cell lines. To determine if daraxonrasib could block adaptive resistance to trametinib and reactivation of ERK, we treated SUM159PT (HRAS-G12D), SUM102PT (wt-RAS), and MDA-MB-468 (wt-RAS) with trametinib or daraxonrasib alone and in combination (Figure 6B). Like trametinib, single agent treatment with daraxonrasib resulted in a transient decrease in activating phosphorylation of ERK after four hours of treatment that partially returned at 48 hours. However, daraxonrasib did not induce hyperphosphorylation of MEK, consistent with inhibition of RAS(on) activation of RAF. Combining the two inhibitors blocked both the hyperphosphorylation of MEK induced by trametinib treatment and the reactivation of phospho-ERK observed with treatment of either compound as a single agent (Figure 6B). These findings collectively support the model that RTK reprogramming induced by trametinib treatment drives activation of RAS and subsequent reactivation of MEK/ERK signaling (Figure 6C). Since daraxonrasib selectively binds RAS-GTP, increased RAS-GTP levels induced by trametinib are inhibited by daraxonrasib preventing RAF activation, and trametinib inhibits MEK phosphorylation of ERK and ERK reactivation.

**Figure 6:**
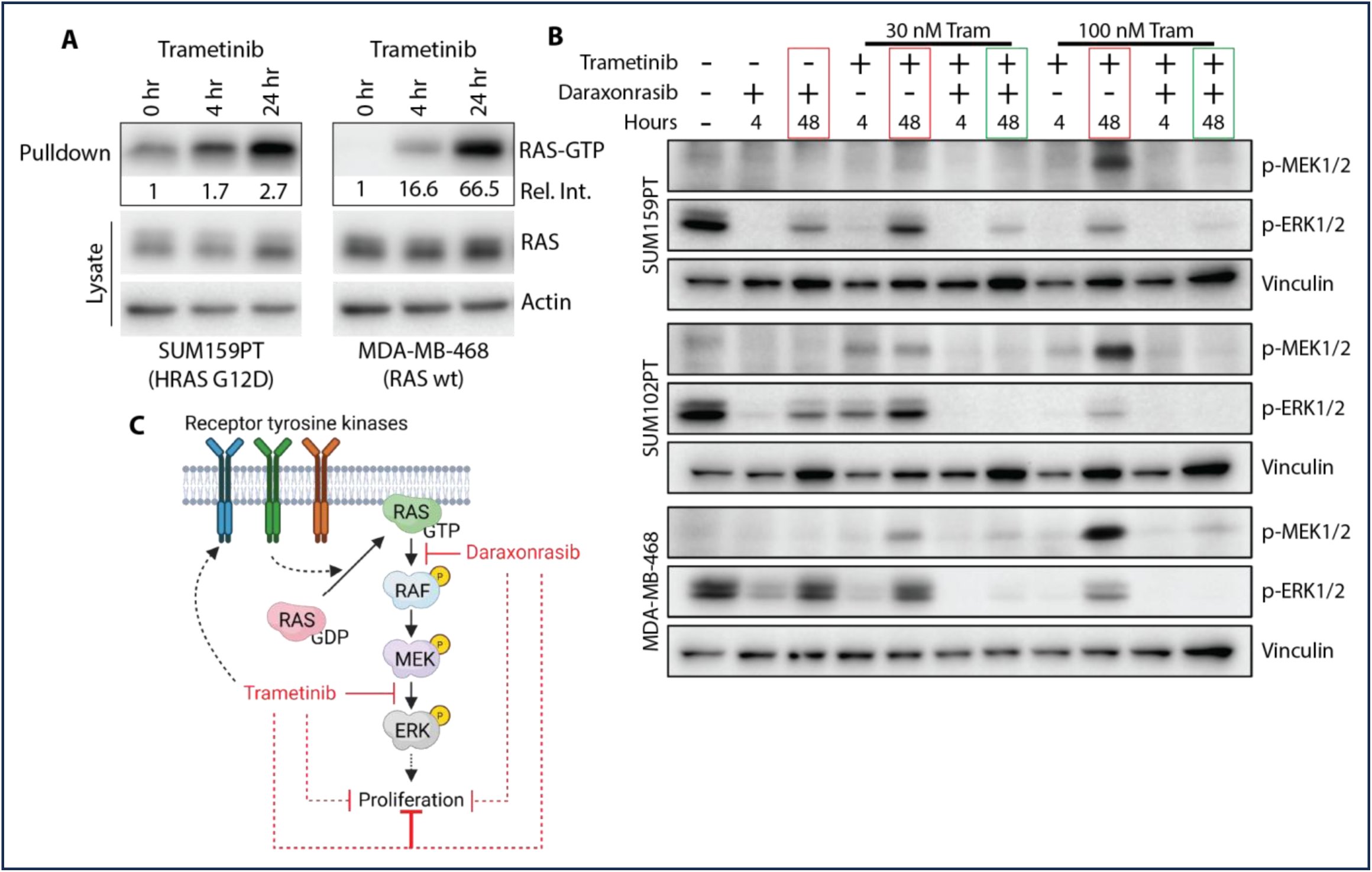
Combining daraxonrasib and trametinib blocks pathway reactivation durably inhibiting MEK/ERK signaling. **A)** Trametinib treatment increased cellular levels of activated RAS. SUM159PT or MDA-MB-468 cells were treated with 100 nM trametinib for 4 or 24 hours then RAS-GTP levels determined using a RAS-GTP effector pulldown assay. The pulldown assay uses the binding domain of cRAF which binds selectively to RAS-GTP conjugated to agarose beads to precipitate activated RAS from cell lysate. **B)** Combinations of trametinib and daraxonrasib blocked MEK/ERK pathway reactivation observed with either single agent treatment. Three TNBC cell lines were treated with 100 nM daraxonrasib and/or two different concentrations of trametinib (30 nM or 100 nM) for 4 or 48 hours. Levels of activating phosphorylation of MEK1/2 and ERK1/2 were measured by immunoblot using phospho-specific antibodies. Vinculin was used as a loading control. Red and green boxes highlight single agent and combination treatment lanes at 48 hours, respectively. **C)** Model describing how daraxonrasib blocks adaptive resistance to trametinib. At early time points following trametinib treatment MEK/ERK signaling is effectively blocked resulting in transient growth arrest. Within 24 hours of trametinib treatment, the RTK landscape is remodeled driving activation of RAS-GTP leading to hyperphosphorylation of MEK and reactivation of ERK to re-establish cell proliferation. By combining daraxonrasib with trametinib, activated RAS-GTP is sequestered preventing activation of RAF kinases, hyperphosphorylation of MEK, and re-activation of ERK and cell proliferation. Panel C was created in BioRender.

### Daraxonrasib and trametinib inhibit tumor growth and increase survival in mice expressing TNBC PDXs

Treatment of TNBC cell lines *in vitro* with daraxonrasib in combination with trametinib durably inhibited the RAS-RAF-MEK-ERK signaling pathway and effectively suppressed cellular proliferation (Figures 5&6). We tested the effects of daraxonrasib and trametinib alone and in combination using WHIM2 and WHIM30 TNBC patient derived xenografts (PDXs) in NSG mice. Tumor volume measurements were taken 14 days after initiation of treatment (Figure 7A and B) and survival response shown in Figures 7C and D for WHIM2 and WHIM30 PDXs, respectively. The two PDXs have different responses to daraxonrasib and trametinib. For the WHIM2 PDX, each inhibitor as a single agent modestly but statistically significantly inhibited tumor growth and survival (Figure 7A and C). Combination daraxonrasib + trametinib treatment strongly inhibited tumor growth and markedly increased survival. In contrast, WHIM30 PDXs were highly responsive to both daraxonrasib and trametinib as single agents (Figure 7B and D). Combination daraxonrasib + trametinib gave a statistically significant increase in survival relative to single agent treatments with the WHIM30 tumors. The results demonstrate that PDXs have a differential sensitivity to daraxonrasib and trametinib but that the combination therapy inhibited tumor growth and significantly increased survival.

**Figure 7:**
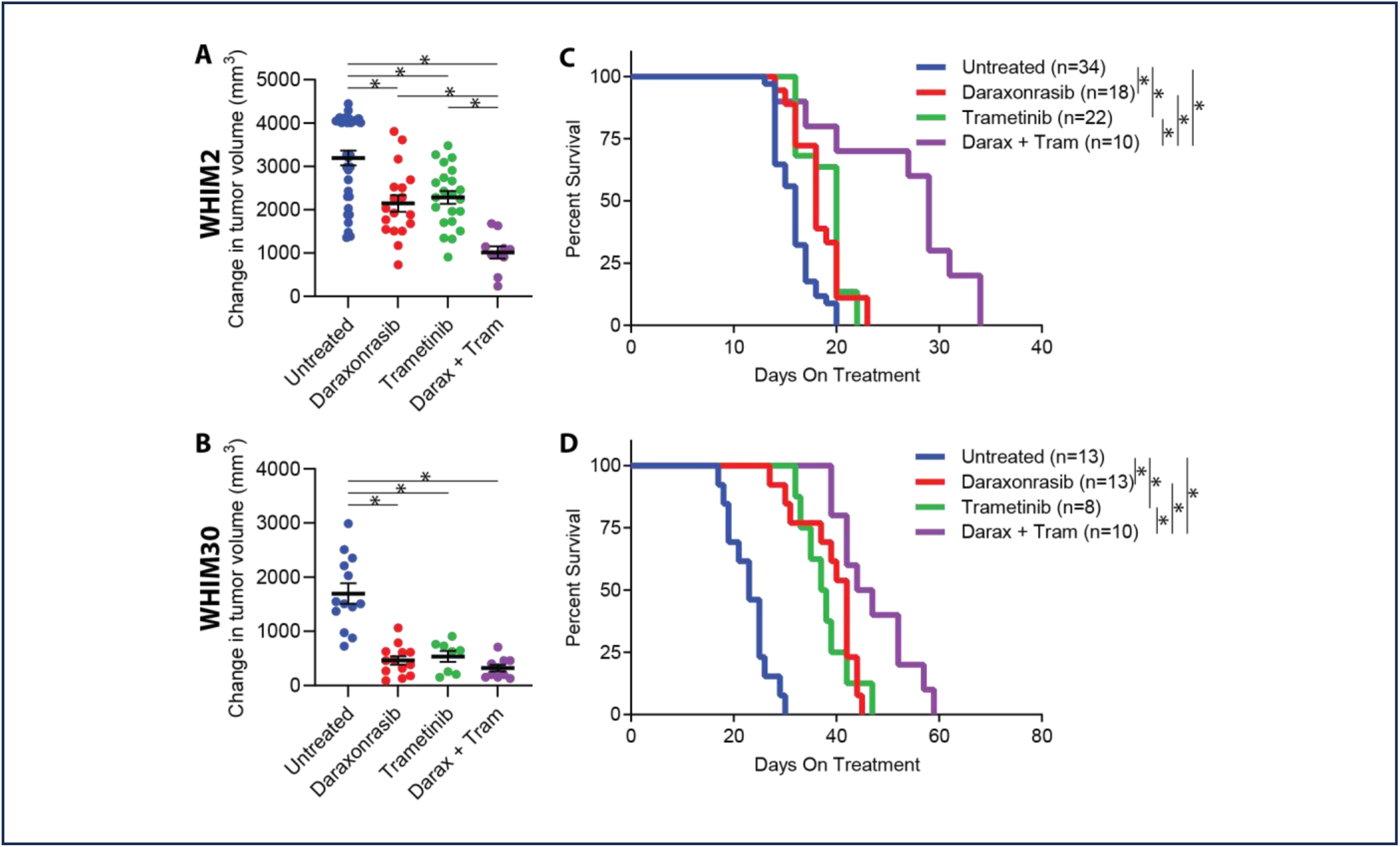
Combining RMC-6236 and trametinib improves survival of TNBC patient derived xenografts *in vivo*. WHIM2 or WHIM30 cells were implanted into the fourth mammary fat pad of NSG mice. Treatment was initiated with 25 mg/kg daraxonrasib and/or 1 mg/kg trametinib daily through chow upon tumors reaching 5-6 mm in size. **A and B)** Tumor volume was measured using calipers after 14 days of treatment. **C and D)** Kaplan-Meier plots show improved survival with either single agent treatment and combination treatment compared to untreated mice. An (*) indicates statistical significance for the indicated comparison (p < 0.05) using unpaired, two-tailed students T-tests.

## Discussion

Kinome analysis of tumors from two clinical trials of TNBC patients treated with the MEK inhibitor trametinib demonstrated that each tumor had a different quantitative expression of RTK expression before and after treatment, overlapping with RTK expression in TNBC organoids and cell lines at baseline and after MEK inhibitor exposure^6, 8^ (Suppl Figure 2). We previously demonstrated that MEK inhibitor-induced adaptive RTK expression results in activating tyrosine phosphorylation of the RTKs (e.g., PDGFRB, VEGFR2, RET) in SUM159PT cells^5^. Exposure of TNBC cells to MEK inhibitor activates RAS GTP binding because of adaptive reprogramming. Thus, heterogeneity of expression of RTKs makes it basically impossible for a single targeted kinase inhibitor or even a combination of inhibitors to have clinical benefit for a spectrum of patients. This conclusion is consistent with the clinical trial using erlotinib as an EGFR inhibitor where only ∼20% of TNBC patient tumors had a growth inhibitory response, correlating with the frequency of EGFR amplification in TNBC tumor cells^26^.

RAS activation of the RAF-MEK-ERK pathway is a predominant regulator of proliferation, with approximately 60% of TNBC having amplified KRAS or its effector kinase BRAF upstream of MEK-ERK with gain of function mutations being rare. The pan-RAS(on) inhibitor daraxonrasib overcomes the variable RTK landscape and amplification of RAS and RAF proteins by stabilizing a RAS(on)-cyclophilin A protein complex unable to bind and activate RAS regulated proteins including RAF kinases. Daraxonrasib, as a single agent, had significant growth inhibitory action *in vitro* with cell lines and *in vivo* with human WHIM2 and WHIM30 PDXs. However, using an *ex vivo* primary human TNBC tumor preparation, daraxonrasib treatment was shown to induce a transcriptional reprogramming response similar to that seen with trametinib inhibition of MEK-ERK involving the expression of different RTKs. Consistent with this adaptive reprogramming and adaptive bypass of MEK inhibition, ERK reactivation was also observed in cell lines treated with daraxonrasib indicating an escape mechanism for single agent inhibitors. Combining daraxonrasib and trametinib overcame adaptive bypass and ERK reactivation and enhanced efficacy *in vitro* and *in vivo* when comparing either treatment alone. Daraxonrasib was effective at nanomolar concentrations in sensitizing cells to trametinib inhibition of cell growth, independent of their kinotype or level of cMYC expression. Our studies indicate that daraxonrasib in combination with trametinib inhibits growth of RTK-driven TNBC cells expressing wild-type RAS or mutant RAS protein.

To our knowledge, daraxonrasib has not been tested before in TNBC and we report efficacy as a single agent or in combination with a MEK inhibitor irrespective of oncogenic RAS mutation. This combination therapy provides a new strategy for the treatment of RTK driven tumors such as TNBC having wild type or mutant RAS proteins, independent of the RTK expression landscape of the tumor. Thus, pan-RAS(on) inhibition alone or in combination is a clinically relevant, innovative therapy for the treatment of TNBC complementary and mechanistically different from standard of care chemotherapy, immunotherapy or antibody-drug conjugates. Daraxonrasib as a single agent or in combination therapy is relevant to many different cancers that are RTK driven having either wild-type or mutant RAS proteins.

## Resource Availability

### Lead contact

Requests for further information and resources should be directed to and will be fulfilled by the lead contact, Gary L. Johnson.

### Materials availability

There are restrictions to the availability of the isotopically labeled heavy peptides used in this study due to the lack of an external centralized repository for its distribution and our need to maintain the stock. Peptide sequences and standard operating procedures for the handling of synthesized peptides are available through the Clinical Proteomic Tumor Analysis Consortium Assay Portal (proteomics.cancer.gov/assay-portal/) and in our supplementary materials.

## Acknowledgements

This work was supported by the UNC Lineberger Triple Negative Breast Cancer (TNBC) Center (GLJ, PMS, CMP), NIH U24DK116204 as part of the Illuminating the Druggable Genome Project (GLJ, RRT, SMG), NCI Breast SPORE P50CA058223 (CMP, HSE, LAC), NCI U01CA238475 (CMP, GLJ), NCI K08CA280388 (PMS), NCI R01CA233811 (SMG), NCI R01CA288145 (JJY, GLJ), NCI U01CA274298 (JJY, GLJ), Pancreatic SPORE P50CA257911 (JJY, GLJ), UNC LCCC Support Grant NCI P30CA016086 (HSE), UNC Lineberger Developmental Funding Program (MPE, DOO), NCI P01CA013106-Project 3 and CSHL/Northwell Health (DLS). We thank the UNC LCCC Tissue Procurement Core for breast cancer tumor specimens, the Translational Genomics Lab core facility (SCR_025231), and the UNC Metabolomics and Proteomics (MAP) Core for support of the Exploris 480 MS. Genomics project management was performed by the Lineberger Office of Genomics Research at the University of North Carolina-Chapel Hill, which is supported in part by the Lineberger University Cancer Research Fund, by the NIH NCI 5UG1CA233333 grant, and The UNC Center for Environmental Health and Susceptibility (UNC-CEHS) P30ES010126 grant. The Washington University Proteomics Shared Resource (WU-PSR) is supported in part by the WU Institute of Clinical and Translational Sciences (NCATS UL1TR000448), the Mass Spectrometry Research Resource (NIGMS P41GM103422) and the Siteman Comprehensive Cancer Center Support Grant (NCI P30CA091842) and CPTAC (U24CA160035). The expert technical assistance of Alan Davis and Rose Connors is gratefully acknowledged. We would like to acknowledge the contributions of Bhuvaneswari Ramaswamy, MD in memoriam.

## Author Contributions

Conceptualization, M.P.E., R.W.S., Q.Z., N.S., M.D.S., C.M.P., S.M.G., R.R.T., and G.L.J.; formal analysis, M.P.E., R.W.S., and F.O.Q.; funding acquisition, M.P.E., D.O.O., C.M.P., H.S.E., L.A.C., J.J.Y., D.L.S., S.M.G., P.M.S., R.R.T., and G.L.J.; investigation, M.P.E., R.W.S., F.O.Q., D.O.O., A.A.W., C.U.J., X.C., P.E.G., Y.M., I.C.M., X.Y., and J.S.Z.; methodology, M.P.E., R.W.S., S.B., D.L.S., and R.R.T. PMS; resources, S.B., P.A.S., M.J.E., H.S.E., L.A.C., J.J.Y., D.L.S., P.M.S., R.R.T., and G.L.J.; software, J.L.; visualization, M.P.E., F.O.Q., and J.P.M.; Writing – original draft, M.P.E. and G.L.J.; writing – review and editing, M.P.E., R.W.S., F.O.Q., D.O.O., Q.Z., S.M.G., P.M.S., R.R.T., and G.L.J.

## Supplemental Information

Document S1. Figures S1 – S2.

Table S1. SIL peptide sequences

## Methods

### Cell culture

Cell lines and organoids were maintained at 37°C, 5% CO_2_. HCC1143, HCC1806, HCC70, Hs578T, SK-BR-3, and BT-474 were grown in RPMI supplemented with 10% fetal bovine serum (FBS) and 1% PennStrep (Gibco). MDA-MB-468 cells were grown in DMEM supplemented with 10% FBS and 1 % PennStrep. SUM102PT and SUM149PT cells were grown in complete HUMEC media supplemented with 10% FBS and 1% PennStrep. SUM159PT and MDA-MB-231 cells were grown in DMEM/F12 supplemented with 5% FBS, 1% PennStrep, 5 ug/mL insulin, and 1 ug/mL hydrocortisone. PDO lines were grown in 300 uL domes of Cultrex Ultimatrix reduced growth factor basement membrane (Bio-Techne), and maintained in DMEM/F12 (Gibco), 10% R-Spondin1 conditioned media obtained from Cultrex® HA-R-Spondin1-Fc 293T Cells (Bio-Techne), B27 supplement (Gibco), 5 mM Nicotinamide (Sigma-Aldrich), 1.25 mM N-Acetylcysteine (Sigma-Aldrich), 50 ug/ml Primocin (InvivoGen), 100 ng/ml Noggin (PEPROTECH), 5 ng/ml hEGF (PEPROTECH), 37.5 ng/ul Heregulin-beta (PEPROTECH) 5 ng/ml FGF-7 (PEPROTECH), 20 ng/ml FGF10 (PEPROTECH), 5uM Y-27632 (Abmole Bioscience), 500 nM SB202190 (Sigma-Aldrich), 500 nM A83-01 (Tocris) as previously described^29^.

### Tumor samples

Patient tumor samples were provided by the UNC Tissue Procurement Core Facility as either formalin fixed, paraffin embedded 10µm scrolls or fresh needle biopsies. All samples were collected with informed written consent from patients pursuant with the University of North Carolina’s Institutional Review Board. Fresh needle biopsies for *ex vivo* culture were immediately minced with sterile razor blades and evenly distributed into separate wells containing DMDM/F12 medium supplemented with 10% FBS and 1% Penn/Strep with vehicle (DMSO) or 100 nM daraxonrasib and cultured for 24 hours at 37C and 5% CO_2_. Both cells in suspension and adherent cells were collected by pipetting and scraping, respectively. Cells were washed with PBS and flash frozen until processing for RNAseq.

### Stable isotope labeled peptides

SIL peptides were purchased from New England Peptides’ custom synthesis service with uniformly C13 and N15 labeled- C-terminal amino acids at > 95% chemical and 99% isotopic purity. SIL peptides were dissolved in a solution containing 30% acetonitrile and 1% formic acid, combined at equimolar ratios, aliquoted for single usage, and lyophilized for storage at -80°C. The day of solid phase extraction of endogenous peptides, SIL peptides were dissolved in 2% acetonitrile and 0.1% formic acid to a final concentration of 25 fmol/µL for sample spike in.

### Sample preparation

For mass spectrometry analysis using SureQuant/PRM, cell line and PDO samples were lysed in buffer containing 50 mM Tris pH 8.0, 8 M urea, 75 mM NaCl, 1 mM EDTA, 2 µg/mL aprotinin, 10 µg/mL leupeptin, 1 mM phenylmethylsufonyl fluoride, 10 mM NaF, and 1% each of Sigma phosphatase inhibitor cocktails 2 and 3. Samples were sonicated using three 10 second bursts using a microprobe sonicator set to 30% amplitude prior to clarification by centrifugation at 16,000 × *g* for 30 minutes at 8°C. Supernatants were transferred to fresh microfuge tubes and the bicinchonic acid assay (BCA) was used to determine protein concentration. A total of 5 µg of protein was reduced using 5 mM DTT for 45 minutes at room temperature followed by alkylation with 10 mM iodoacetamide at room temperature in the dark for 45 minutes. Urea was diluted 5-fold using 50 mM Tris pH 8.0 prior to addition of 0.1 µg of Lyc-C. Samples were incubated at room temperature for two hours prior to addition of 0.1 µg of trypsin for overnight digestion at room temperature. Samples were quenched by addition of trifluoroacetic acid (TFA) to pH < 2.0. SIL peptides were added at the defined ratio of 25 fmol per microgram of endogenous protein. Samples with SIL spike in were then desalted using Pierce peptide desalting columns according to manufacturer instructions. Eluted peptides were lyophilized and stored at -80°C until LC-MS/MS analysis.

For FFPE samples, 10 µm scrolls were deparaffinized using three five-minute xylene incubations. Deparaffinized samples were then rehydrated using sequential washes consisting of 100%, 85%, and 70% ethanol. Rehydrated pellets were then resuspended in 100 mM ammonium bicarbonate, pH 8.0 prior to incubation at 80°C to reverse crosslinking. Samples were cooled to room temperature prior to addition of 2,2,2 trifluoroethanol to a final concentration of 50%. Samples were sonicated for 25 seconds in five second pulses using a microtip sonicator set to 30% amplitude then incubated at 60°C for an hour. Sonication and incubation at 60°C was repeated one additional time. Protein concentration was determined using the BCA assay followed by reduction using 13.3 mM Tris(2-carboxyethyl)phosphine and 33.3 mM DTT for 30 minutes at 60°C. Samples were alkylated using 50 mM iodoacetamide for 20 minutes in the dark at room temperature. Samples were further diluted by 2.5-fold using 50 mM ammonium bicarbonate, pH 8.0 prior to addition of trypsin at a w:w ratio of 1:50 trypsin:total protein. Proteolysis was quenched by adding TFA so pH < 2.0 and peptides desalted using Pierce peptide desalting columns according to manufacturer instructions. Eluants were lyophilized and resuspended in 2% acetonitrile and 0.1% formic acid for quantitation of total peptide using the Pierce quantitative fluorometric peptide assay. SIL peptides were added at a final ratio of 25fmol SIL peptide per microgram of total peptide. Samples were then lyophilized and stored at -80°C until LC-MS/MS analysis.

### LC-MS/MS analysis

Proteomics samples were analyzed using a Thermo Orbitrap Exploris 480 mass spectrometer with a Nanospray Flex ion source with Sonation column oven and an UltiMate 3000 HPLC. Peptides were separated using a 15 cm Aurora Elite column (IonOpticks) heated to 40°C. Mobile Phase A (MPA) consisted of 0.1% formic acid in water with Mobile Phase B (MPB) as 80% acetonitrile and 0.1% formic acid. The elution gradient consisted of sequential linear gradients from 3-19% MPB over 72 minutes, 19-29% MPB over 28 minutes, 29-41% MPB over 20 minutes, and 41-95% MPB over 3 minutes followed by 7 minutes of constant 95% MPB. The flow rate for the elution gradient was 250 nL/min. MS settings include 2.1 kV spray voltage, 275°C ion transfer tube temperature. Full scan settings were a scan range of 300 – 1500, resolution of 120,000, normalized AGC target of 300%, an inject time of 50 ms, a 3 s cycle time, and +/- 3 ppm mass tolerance. Upon detection of a matching precursor ion during MS1, a 7,500 resolution MS2 scan for the SIL peptide was triggered using 150 – 1700 scan range, normalized AGC target of 1000%, injection time of 10 ms, and HCD collision energy of 27%. Pseudomatching of at least 3 product ions was required to trigger a 60,000 resolution scan for endogenous peptide fragments with m/z offsets specific for the isotopic label using 150 – 1700 scan range, 1000% normalized AGC target, inject time of 90 ms, and HCD collision energy of 27%.

### Mass spectrometry data analysis

SureQuant/PRM data were analyzed using the Skyline software package^30, 31^. Ratios of interpolated peak areas were calculated for the SIL peptides compared to the endogenous peptide using the three most abundant product ions and requiring at least six points across peaks. Specificity was assessed by qualitative analysis of peak shape, retention time, and the relative distribution of all product ions and a mass error threshold of approximately 10 ppm. When three product ions were not available, two product ions were used for quantitation. Peptide quantitation with > 30% coefficients of variance from biological replicates were discarded for cell lines and organoid samples. A single peptide was used for protein quantitation selected based on the most abundant quantitation that was free from interference or other confounding factors. Kinome trees were generating using the CORAL package^32^.

### RNAseq

Samples processed for RNAseq were collected by scraping in ice-cold PBS and pelleting by centrifugation for cell lines or by grinding tumor samples using a mortar and pestle. RNA was isolated using the Qiagen RNeasy Plus kit per the manufacturer’s instructions. cDNA libraries were prepared using the Roche Kapa Hyperprep kit according to the manufacturer’s instructions and paired end sequencing performed on the Illumina Nextseq 1000 sequencing platform. FASTQ files were aligned to the hg38/GRCh38 reference genome using GENCODE v36 annotations using STAR with quantitation performed using Salmon. When comparing baseline expression levels across samples, Salmon abundances were upper quartile normalized. Differential expression analysis was performed using DESeq2 after rounding Salmon abundances to the nearest integer. Statistical significance threshold was set at p-adj < 0.05 for each pairwise analysis.

### Immunoblotting and RAS-GTP pulldown assay

For immunoblot analysis, cells were washed with ice-cold PBS and harvested by scraping. Cells were pelleted at 500 × *g* at 4°C for five minutes. After removing the supernatant, cells were lysed in RIPA buffer consisting of 10 mM Tris-HCl pH 7.6, 140 mM NaCl, 1 mM EDTA, 0.5 mM EGTA 1% Triton X-100, 0.1% sodium deoxycholate, 0.1% SDS, 10 mM NaF, 2.5 mM NaVO_4_, Phosphatase Cocktail 2 (Sigma), Phosphatase Cocktail 3 (Sigma) and Protease Inhibitor Cocktail (Roche). Lysates were incubated on ice for 10 minutes, sonicated three times for 10 seconds at 20% amplitude using a microprobe sonicator, and clarified by centrifugation at 16,000 × *g* at 4°C for 15 minutes. Protein content was determined using Thermo BCA assay. Proteins were resolved by SDS-PAGE and transferred to nitrocellulose membranes. Membranes were blocked for one hour in 5% powdered milk in tris buffered saline containing 0.1% tween-20 (TBST) then with primary antibody in 5% BSA in at 4°C overnight. Membranes were washed for 10 minutes in TBST three times then incubated with HRP-conjugated secondary antibody in 5% milk in TBST. Membranes were washed three more times and imaged with a ChemiDoc imager (BioRad). Primary antibodies: pan-RAS (DSHB, CPTC-KRAS4B-2-s), pERK1/2-T202/Y204 (Cell Signaling Technologies (CST), 4370), pMEK-S217/S221 (CST, 9121), vinculin (CST, 13901), β-actin (Santa Cruz Biotechnology (SCBT), sc-47778), ARAF(SCBT, sc-408), BRAF (SCBT, sc-166), CRAF (SCBT, sc-227), HRAS (proteintech, 18295-1-AP), KRAS (SCBT, sc-30), NRAS (SCBT, sc-31), c-MYC (Abcam, ab32072).

RAS activation was measured using the active RAS detection kit (Cell Signaling Technology) according to manufacturer recommendations. Briefly, cells were washed with ice-cold PBS and scraped directly into the provided lysis buffer. Lysates were incubated on ice for 5 minutes then clarified by centrifugation at 16,000 × *g* at 4°C for 15 minutes. 1,000 µg of lysate protein per sample was incubated with 100 µL (80 µg) of Raf-RBD bead slurry for 1 hour at 4°C with end-over-end rotation. Beads were washed once with lysis buffer and bound proteins eluted with 2X Laemmli buffer (125 mM Tris pH 6.8, 20% glycerol, 4% SDS, 0.005% bromophenol blue, and 5% β-mercaptoethanol).

### Viability and colony formation assays

For 96-hour growth assays, cells were seeded at low density on to 96-well plates with clear bottoms and opaque walls. Media was exchanged the following day with complete growth media containing the indicated concentration of drug(s). Cells were incubated at 37°C and 5% CO_2_ for 96 hours when CellTiter-Glo (Promega) reagent was added directly to the growth media. Plates were incubated at room temperature for 20 minutes then luminescence measured using a PheraStar Microplate Reader (BMG). Growth curves were generated based on drug treated luminescence relative to vehicle control.

For colony formation assays, cells were seeded at low density onto 12-well plates. The following day, media was exchanged for complete growth media containing the indicated concentration of drug(s). Plates were incubated with drug at 37°C and 5% CO_2_ for a total of 14 days with media/drug replaced every three days. Cells were then washed once with ice-cold PBS and incubated with staining buffer consisting of 0.5% crystal violet and 20% methanol at room temperature for 15 minutes. Plates were then washed by submersion in water three times or until water washes were free of coloration. Plates were dried at room temperature prior to imaging. Crystal violet was then solubilized in buffer containing 100 mM sodium citrate pH 4.2 and 50% ethanol. Absorption of the dissolved crystal violet was determined using a PolarStar microplate reader (BMG) at a wavelength of 570 nm.

### Mouse studies

All animal work was conducted in accordance with the Institutional Animal Care and Use Committee (IACUC) guidelines. Only female mice were used for all experiments without blinding. PDX transplantation was performed as described previously^33, 34^. Briefly, tumors were digested to form cell aggregate suspensions which were washed in Hank’s Balanced Salt Solution containing 2% FBS and resuspended in the same media with 50% Matrigel. Mice were anesthetized with 2% isoflurane and tumor cells injected in the fourth mammary fat pad of NSG mice. Mice were monitored 2-3 times per week for tumor growth using caliper measurements. Upon reaching 5 mm in any dimension, mice were randomized into treatment groups and treatment was initiated with 1 mg/kg trametinib, 25 mg/kg daraxonrasib, or a combination of both inhibitors daily through chow. Pre-treatment body weight was recorded for each mouse and mice were closely monitored for weight loss and body condition score (BCS) every 2 – 3 days as well as for lethargy or aberrant behavior. Animals were humanely euthanized upon tumors reaching 20 mm in any dimension. Additional criteria for euthanasia were set including ≥ 20% loss in body weight, a BCS of 2 or below, or formation of multiple tumors were applied but no mice met any of these criteria. Tumor volume was calculated using the formula volume = (L x W^2^)/2, where L and W were tumor length and width, respectively, measured in millimeters.

---

**Supplementary Figure 1.**
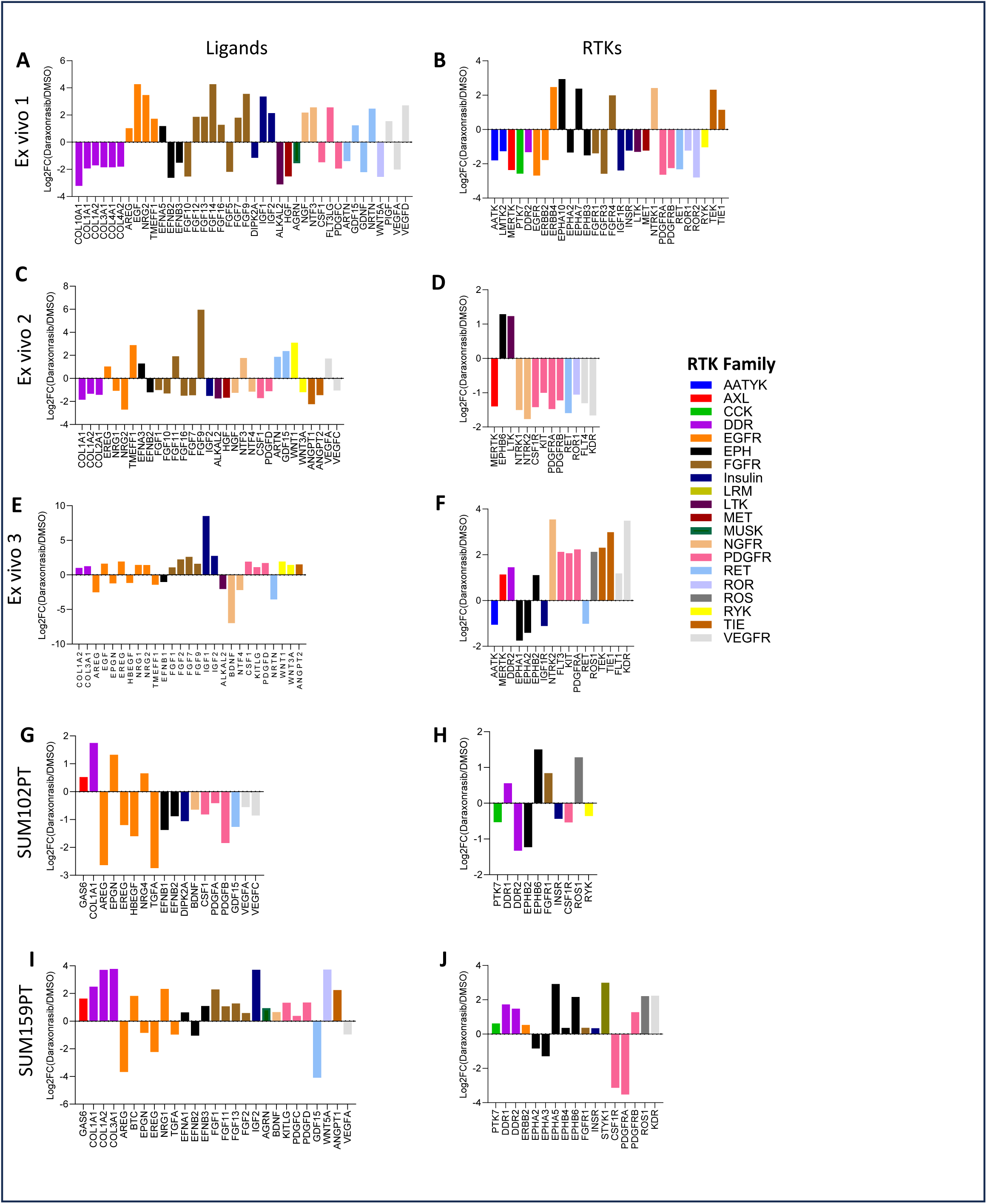
: Daraxonrasib induced dynamic reprogramming of the RTK and RTK ligand landscape in *ex vivo* patient tumor samples and cell lines. Cell lines or *ex vivo* cultures were treated with 100 nM daraxonrasib for 24 hours prior to RNAseq analysis. Plotted are genes that were differentially expressed (DEGs) in response to treatment. Differential expression was defined as at least a 2-fold change in expression for *ex vivo* cultures where biological replicates were not possible (A-F). Differential gene expression was defined by padj < 0.05 following DEseq2 in cell lines where treatments were performed in biological triplicate (G-J). Expression changes in ligands are shown in the left side panels and RTKs in the right side panels. Ligands and RTKs were color-coded based on their respective RTK family^27, 28^.

**Supplementary Figure 2.**
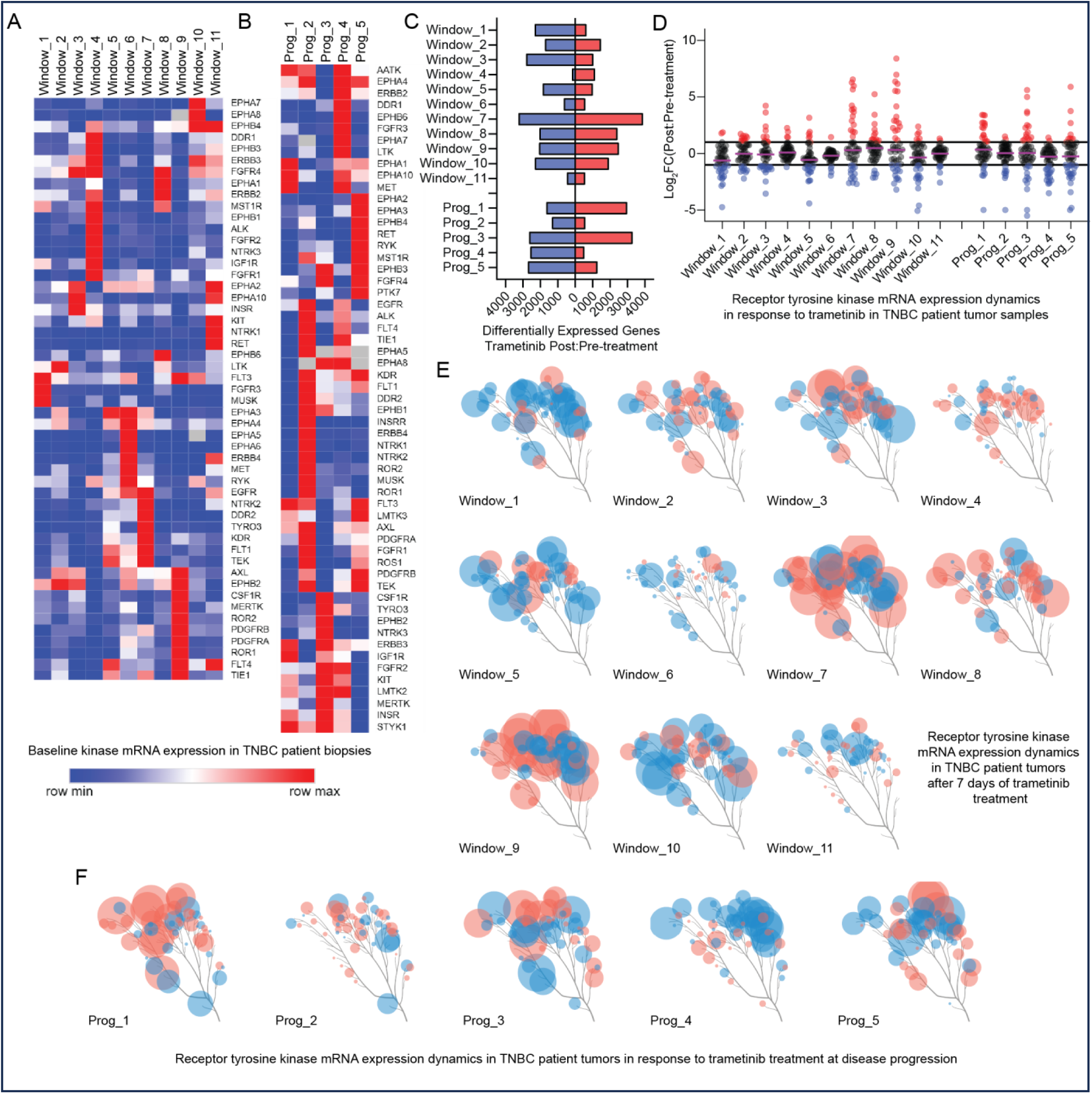
: Dynamic variability in expression of receptor tyrosine kinases in TNBC patient tumors. The effects of trametinib inhibition of the MEK-ERK pathway on expression of RTKs were determined in two TNBC clinical trials: 1) A window-of-opportunity trial in which untreated TNBC patients awaiting surgery received trametinib daily for seven days. Pre-treatment needle biopsies were taken prior to initiation of trametinib treatment and post-treatment samples taken during surgical resection of the tumor^6^. 2) A phase II clinical trial designed to test the response of trametinib as a single agent and in combination with the pan-AKT inhibitor uprosertib upon reaching progression with trametinib monotherapy in metastatic TNBC patients who previously received chemotherapy^8^. Needle biopsies were taken pre-treatment and upon progression with single agent trametinib treatment which ranged from 2.6 to 5.6 months. Tumor samples for both trials were analyzed via RNAseq. **A and B)** Distinct RTKs were enriched in each patient tumor in pre-treatment samples from both the window trial (**A**) and progression trial (**B**). Heatmaps indicate the relative expression levels of quantile normalized read counts. **C-F)** For each patient, the post:pre treatment change in transcript abundance for each gene was calculated using quantile normalized read counts. **C)** The number of differentially expressed genes (DEGs) are summarized for each patient. DEGs were defined as transcripts whose abundance changed by at least 2-fold in response to trametinib treatment. Red and blue bars indicate transcripts that were up or down regulated by trametinib treatment, respectively. **D)** Dynamic reprogramming of RTK transcripts is summarized for each patient. Each circle represents an expressed RTK. **E and F)** RTK kinome trees summarize the dynamic changes RTK transcripts in response to trametinib treatment for window trial (E) and progression trial (F) patients. Each circle indicates an RTK that was expressed in the indicated patient. Red and blue colors indicate up and down regulated transcripts, respectively. Circle size correlates with the magnitude of change with the largest circles indicating an 8-fold or larger change in expression.

## References

1. Leon-Ferre, R. A. & Goetz, M. P. Advances in systemic therapies for triple negative breast cancer. BMJ 381, e071674 10.1136/bmj-2022-071674 (2023).

2. Schmid, P., et al. Pembrolizumab for Early Triple-Negative Breast Cancer. N Engl J Med 382, 810–821 10.1056/NEJMoa1910549 (2020).

3. Cortes, J., et al. Pembrolizumab plus chemotherapy versus placebo plus chemotherapy for previously untreated locally recurrent inoperable or metastatic triple-negative breast cancer (KEYNOTE-355): a randomised, placebo-controlled, double-blind, phase 3 clinical trial. Lancet 396, 1817–1828 10.1016/S0140-6736(20)32531-9 (2020).

4. Cancer Genome Atlas, N. Comprehensive molecular portraits of human breast tumours. Nature 490, 61–70 10.1038/nature11412 (2012).

5. Duncan, J. S., et al. Dynamic reprogramming of the kinome in response to targeted MEK inhibition in triple-negative breast cancer. Cell 149, 307–321 10.1016/j.cell.2012.02.053 (2012).

6. Zawistowski, J. S., et al. Enhancer Remodeling during Adaptive Bypass to MEK Inhibition Is Attenuated by Pharmacologic Targeting of the P-TEFb Complex. Cancer Discov 7, 302–321 10.1158/2159-8290.CD-16-0653 (2017).

7. Goulet, D. R., et al. Discrete Adaptive Responses to MEK Inhibitor in Subpopulations of Triple-Negative Breast Cancer. Mol Cancer Res 18, 1685–1698 10.1158/1541-7786.MCR-19-1011 (2020).

8. Prasath, V., et al. Phase II study of MEK inhibitor trametinib alone and in combination with AKT inhibitor GSK2141795/uprosertib in patients with metastatic triple negative breast cancer. Breast Cancer Res Treat 210, 179–189 10.1007/s10549-024-07551-z (2025).

9. Graves, L. M., Duncan, J. S., Whittle, M. C. & Johnson, G. L. The dynamic nature of the kinome. Biochem J 450, 1–8 10.1042/BJ20121456 (2013).

10. East, M. P. & Johnson, G. L. Adaptive chromatin remodeling and transcriptional changes of the functional kinome in tumor cells in response to targeted kinase inhibition. J Biol Chem 298, 101525 10.1016/j.jbc.2021.101525 (2022).

11. Xue, J. Y., et al. Rapid non-uniform adaptation to conformation-specific KRAS(G12C) inhibition. Nature 577, 421–425 10.1038/s41586-019-1884-x (2020).

12. Klomp, J. E., Klomp, J. A. & Der, C. J. The ERK mitogen-activated protein kinase signaling network: the final frontier in RAS signal transduction. Biochem Soc Trans 49, 253–267 10.1042/BST20200507 (2021).

13. Holderfield, M., et al. Concurrent inhibition of oncogenic and wild-type RAS-GTP for cancer therapy. Nature 629, 919–926 10.1038/s41586-024-07205-6 (2024).

14. East, M. P., et al. Quantitative proteomic mass spectrometry of protein kinases to determine dynamic heterogeneity of the human kinome. bioRxiv, 10.1101/2024.10.04.614143 (2024).

15. Manning, G., Whyte, D. B., Martinez, R., Hunter, T. & Sudarsanam, S. The protein kinase complement of the human genome. Science 298, 1912–1934 10.1126/science.1075762 (2002).

16. Jiang, J., et al. Translational and Therapeutic Evaluation of RAS-GTP Inhibition by RMC-6236 in RAS-Driven Cancers. Cancer Discov 14, 994–1017 10.1158/2159-8290.CD-24-0027 (2024).

17. Peterson, A. C., Russell, J. D., Bailey, D. J., Westphall, M. S. & Coon, J. J. Parallel reaction monitoring for high resolution and high mass accuracy quantitative, targeted proteomics. Mol Cell Proteomics 11, 1475–1488 10.1074/mcp.O112.020131 (2012).

18. Gallien, S., Kim, S. Y. & Domon, B. Large-Scale Targeted Proteomics Using Internal Standard Triggered-Parallel Reaction Monitoring (IS-PRM). Mol Cell Proteomics 14, 1630–1644 10.1074/mcp.O114.043968 (2015).

19. Cregg, J., et al. Discovery of Daraxonrasib (RMC-6236), a Potent and Orally Bioavailable RAS(ON) Multi-selective, Noncovalent Tri-complex Inhibitor for the Treatment of Patients with Multiple RAS-Addicted Cancers. J Med Chem 68, 6064–6083 10.1021/acs.jmedchem.4c02314 (2025).

20. Zikry, T. M., et al. Cell cycle plasticity underlies fractional resistance to palbociclib in ER+/HER2- breast tumor cells. Proc Natl Acad Sci U S A 121, e2309261121 10.1073/pnas.2309261121 (2024).

21. Kim, H., et al. Tamoxifen Response at Single-Cell Resolution in Estrogen Receptor-Positive Primary Human Breast Tumors. Clin Cancer Res 29, 4894–4907 10.1158/1078-0432.CCR-23-1248 (2023).

22. Whitman, A. A., et al. Kinase Plasticity with Vandetanib Treatment Enhances Sensitivity to Tamoxifen in Estrogen Receptor Positive Breast Cancer. Mol Cancer Ther, 10.1158/1535-7163.MCT-26-0063 (2026).

23. Klomp, J. E., et al. Determining the ERK-regulated phosphoproteome driving KRAS-mutant cancer. Science 384, eadk0850 10.1126/science.adk0850 (2024).

24. Klomp, J. A., et al. Defining the KRAS- and ERK-dependent transcriptome in KRAS-mutant cancers. Science 384, eadk0775 10.1126/science.adk0775 (2024).

25. Demidenko, E. & Miller, T. W. Statistical determination of synergy based on Bliss definition of drugs independence. PLoS One 14, e0224137 10.1371/journal.pone.0224137 (2019).

26. Gelmon, K., et al. Targeting triple-negative breast cancer: optimising therapeutic outcomes. Ann Oncol 23, 2223–2234 10.1093/annonc/mds067 (2012).

27. Lemmon, M. A. & Schlessinger, J. Cell signaling by receptor tyrosine kinases. Cell 141, 1117–1134 10.1016/j.cell.2010.06.011 (2010).

28. Trenker, R. & Jura, N. Receptor tyrosine kinase activation: From the ligand perspective. Curr Opin Cell Biol 63, 174–185 10.1016/j.ceb.2020.01.016 (2020).

29. Bhatia, S., et al. Patient-Derived Triple-Negative Breast Cancer Organoids Provide Robust Model Systems That Recapitulate Tumor Intrinsic Characteristics. Cancer Res 82, 1174–1192 10.1158/0008-5472.CAN-21-2807 (2022).

30. MacLean, B., *et al*. Skyline: an open source document editor for creating and analyzing targeted proteomics experiments. Bioinformatics 26, 966–968 10.1093/bioinformatics/btq054 (2010).

31. Pino, L. K., et al. The Skyline ecosystem: Informatics for quantitative mass spectrometry proteomics. Mass Spectrom Rev 39, 229–244 10.1002/mas.21540 (2020).

32. Metz, K. S., et al. Coral: Clear and Customizable Visualization of Human Kinome Data. Cell Syst 7, 347–350 e341 10.1016/j.cels.2018.07.001 (2018).

33. Liao, C., et al. Integrated Metabolic Profiling and Transcriptional Analysis Reveals Therapeutic Modalities for Targeting Rapidly Proliferating Breast Cancers. Cancer Res 82, 665–680 10.1158/0008-5472.CAN-21-2745 (2022).

34. Garcia-Recio, S., et al. FGFR4 regulates tumor subtype differentiation in luminal breast cancer and metastatic disease. J Clin Invest 130, 4871–4887 10.1172/JCI130323 (2020).

